# Repetitive blast promotes chronic aversion to neutral cues encountered in the peri-blast environment

**DOI:** 10.1101/2020.02.11.935718

**Authors:** Abigail G. Schindler, Garth E. Terry, Tami Wolden-Hanson, Marcella Cline, Michael Park, Janet Lee, Mayumi Yagi, James S. Meabon, Elaine R. Peskind, Murray M. Raskind, Paul E.M. Phillips, David G. Cook

## Abstract

Repetitive mild traumatic brain injury (mTBI) has been called the “signature injury” of military Servicemembers in the Iraq and Afghanistan wars and is highly comorbid with posttraumatic stress disorder (PTSD). Correct attribution of adverse blast-induced mTBI and/or PTSD remains challenging. Preclinical research using animal models can provide important insight into the mechanisms by which blast produces injury and dysfunction—but only to the degree by which such models reflect the human experience. Avoidance of trauma reminders is a hallmark of PTSD, here we sought to understand whether a mouse model of blast reproduces this phenomenon, in addition to blast-induced physical injuries. Drawing upon well-established work from the chronic stress and Pavlovian conditioning literature, we hypothesized that, even while anesthetized during blast exposure, environmental cues encountered in the peri-blast environment could be conditioned to evoke aversion/dysphoria and reexperiencing of traumatic stress. Using a pneumatic shock tube that recapitulates battlefield-relevant open-field blast forces, we provide direct evidence that stress is inherent to repetitive blast exposure, resulting in chronic aversive/dysphoric-like responses to previous blast-paired cues. The results in this report demonstrate that, while both single and repetitive blast exposures produce acute stress responses (weight loss, corticosterone increase), only repetitive blast exposure also results in co-occurring aversive/dysphoric-like stress responses. These results extend appreciation of the highly complex nature of repetitive blast exposure; and lend further support for the potential translational relevance of animal modeling approaches currently used by multiple laboratories aimed at elucidating the mechanisms (both molecular and behavioral) of repetitive blast exposure.

## INTRODUCTION

Traumatic brain injury (TBI) is a major cause of death and disability, affecting every segment of the population, with youth, elderly, and athletes being most affected. Likewise, following a traumatic event, post-traumatic stress disorder (PTSD) is common and affects 5-10% of the adult population of United States. Moreover, mild traumatic brain injury (mTBI/concussion) is considered the “signature insult” of military Servicemembers of Operation Iraqi Freedom/Enduring Freedom/New Dawn (OIF/OEF/OND) (1, 2), is highly comorbid with posttraumatic stress disorder (PTSD), and is a major source of morbidity among Veterans enrolled in the VA health care system. Such symptoms are common, with approximately 350,000 Veteran mTBIs diagnosed since 2000 (estimated post-deployment rates of 10-25%), with an estimated PTSD comorbidity rate of 50-75% (2-4). Blast exposure caused by detonation of high explosives is the primary source of mTBI (accounting for 75% of all TBIs reported by Veterans) with multiple exposures more common than a single blast exposure (2, 5). Efforts to elucidate the biological processes responsible for the clinical manifestations of blast-related mTBI and PTSD are impeded by several factors that include: (i) high rates of comorbidity; (ii) overlapping symptoms; and (iii) the initiating insult (i.e. blast exposure) can simultaneously induce both biomechanical/neurologic injuries and substantial neuropsychological stress (6, 7). Indeed, clinicians have wrestled with this medical conundrum for more than 100 years since the initial descriptions of “shell shock” among World War I soldiers exposed to artillery bombardments (8, 9). There are at least two competing hypotheses that address the apparent connection between mTBI and PTSD following blast exposure: 1) blast exposure is both a TBI-causing event and a PTSD-related stressor or 2) TBI-induced compensatory changes in stress-related brain regions produce outcomes similar in nature to those seen in PTSD and/or render the brain more susceptible to subsequent stressors (or vice versa – PTSD-related trauma damages brain regions involved in post-concussive symptom outcomes). Following mTBI exposure in rodents, we and others have published evidence of post-concussive syndrome-like and PTSD-like outcomes in both rats and mice (10-15). Anesthesia is commonly used in these preclinical studies. In these settings, the origin of the precipitating blast-induced trauma (physical injury and/or psychological stress) while animals are anesthetized has been viewed alternately as a complicating factor or as a useful reductionistic tool in working toward a better understanding of the mechanisms by which blast exposure results in chronic PTSD-related symptoms (7, 15). While physical injury and neurological damage can result in acute activation of stress pathways, whether blast exposure produces a resulting chronic affective representation of that stress (e.g. PTSD-like aversion/dysphoria to trauma reminders) remains understudied. Translational research efforts using rodent models can provide much needed insight into underlying mechanisms by which blast exposure produces dysfunction, but only in so well as these models recapitulate both battlefield exposure conditions and subsequent injuries and dysfunction displayed by individuals with blast-related mTBI. In this report we sought to understand the extent to which rodent models of blast exposure reproduce the phenomenon experienced by military Servicemembers and Veterans with comorbid mTBI and PTSD.

Drawing upon fundamental concepts regarding chronic stress and Pavlovian fear learning (16-22), we postulated that despite the animals being anesthetized during the blast exposure itself, stress responses attending to events before and after the blast exposure are sufficient to elicit lasting cognitive representations of trauma exposure. We further postulated that such stress responses induce a negative hedonic state with motivational properties that can be associated with neutral cues (odors, visual patterns, etc.) to engender subsequent avoidance and/or reexperiencing-induced stress and dysphoria to those associated cues (16, 17, 19, 22-24). To test these ideas, we examined acute physiological and behavioral stress responses to blast exposure and used place conditioning to measure avoidance and examine behavioral (locomotion and vocalizations) and physiological (corticosterone production) stress responses provoked by re-exposure to blast-related cues. Ultrasonic vocalizations (USVs) are thought to vary with behavioral context, with lower kHz USVs primarily seen during aversive scenarios such as restraint stress or social isolation (25-27), thereby providing a sensitive readout of the animals’ affective state.

Using a well-established electronically-controlled pneumatic shock tube that models battlefield-relevant open-field blast forces generated by detonation of high explosives (10, 11, 28-30), we found that while both single and repetitive blast exposure result in acute stress responses, only repetitive blast exposure produces chronic aversion and dysphoria to prior blast-paired environmental cues. These results offer new insight regarding how repetitive blast exposure may give rise to PTSD-like symptoms. In addition, these findings support the idea that this animal model simultaneously provokes both neurologic and psychological insults, as in military Servicemembers with blast-related comorbid mTBI and PTSD, thus demonstrating a previously unappreciated translational aspect of repetitive blast trauma in rodent models.

## MATERIALS AND METHODS

### Animals and mouse model of blast overpressure

Male C57Bl/6 mice (Jackson Laboratory) aged 3–4 months (weight 22-35 g; mean 27.0 ± 0.2 g) were used. All animal experiments were carried out in accordance with Association for Assessment and Accreditation of Laboratory Animal Care guidelines and were approved by the VA Puget Sound Institutional Animal Care and Use Committees. The shock tube (Baker Engineering and Risk Consultants, San Antonio, TX) was designed to generate blast overpressures that mimic open field high explosive detonations encountered by military Servicemembers in combat, and the design and modeling characteristics have been described in detail elsewhere (10, 28, 29). Briefly, mice were anesthetized with isoflurane (induced at 5% and maintained at 2-3%), secured on a gurney, and placed in the shock tube oriented perpendicular to the oncoming blast wave (ventral body surface facing the oncoming shock wave) (31). Sham (control) animals received anesthesia only for a duration matched to blast animals. Repeated blast/sham exposures occurred successively over the course of three days (one per day). Following exposure, mice were immediately removed from the shock tube and anesthesia was discontinued (anesthesia duration ranged from 3-4 min). Mice were then placed in a warm enclosure for observation during recovery. The blast overpressure (BOP) peak intensity (psi), initial pulse duration (ms), and impulse (psi▪ms) used were in keeping with mild to moderate blast exposure (31, 32) (20.23 psi +/- 0.08 psi; 5.797 ms +/- 0.017 ms; 0.037 psi*ms +/- 0.000 psi*ms) (Figure 1A). Under these experimental conditions, the survival rate was 96%, with blast-exposed mice appearing comparable to sham-exposed mice by inspection 2-4 hours-post blast exposure as previously reported (10, 11, 28, 29). Animals were weighed daily prior to sham/blast exposure and at 24- and 72-hours post-exposure. There were no statistically significant weight differences between 1x sham (n = 9) and 3x sham (n = 18)-treated mice (24h post-exposure: Student’s unpaired t-test, t[25]=0.355, *p*>0.05;. 72h post-exposure: Student’s unpaired t-test, t[25]=0.9368, *p*>0.05), thus, 1x and 3x sham animals at each time point were pooled together for subsequent weight analyses. A subset of animals was sacrificed 30 minutes following the last exposure and trunk blood was collected. Serum samples were processed to assay corticosterone levels using an ELISA kit as per manufacture protocol (Arbor Assays, Ann Arbor, MI). There were no statistically significant corticosterone differences between 1x sham (n = 6) and 3x sham (n = 6)-treated mice (Student’s unpaired t-test, t[10]=0.355, *p*>0.05;. 72h post-exposure: Student’s unpaired t-test, t[25]=0.836, *p*>0.05), thus, 1x and 3x sham animals were pooled together for subsequent corticosterone analyses.

**Figure 1:**
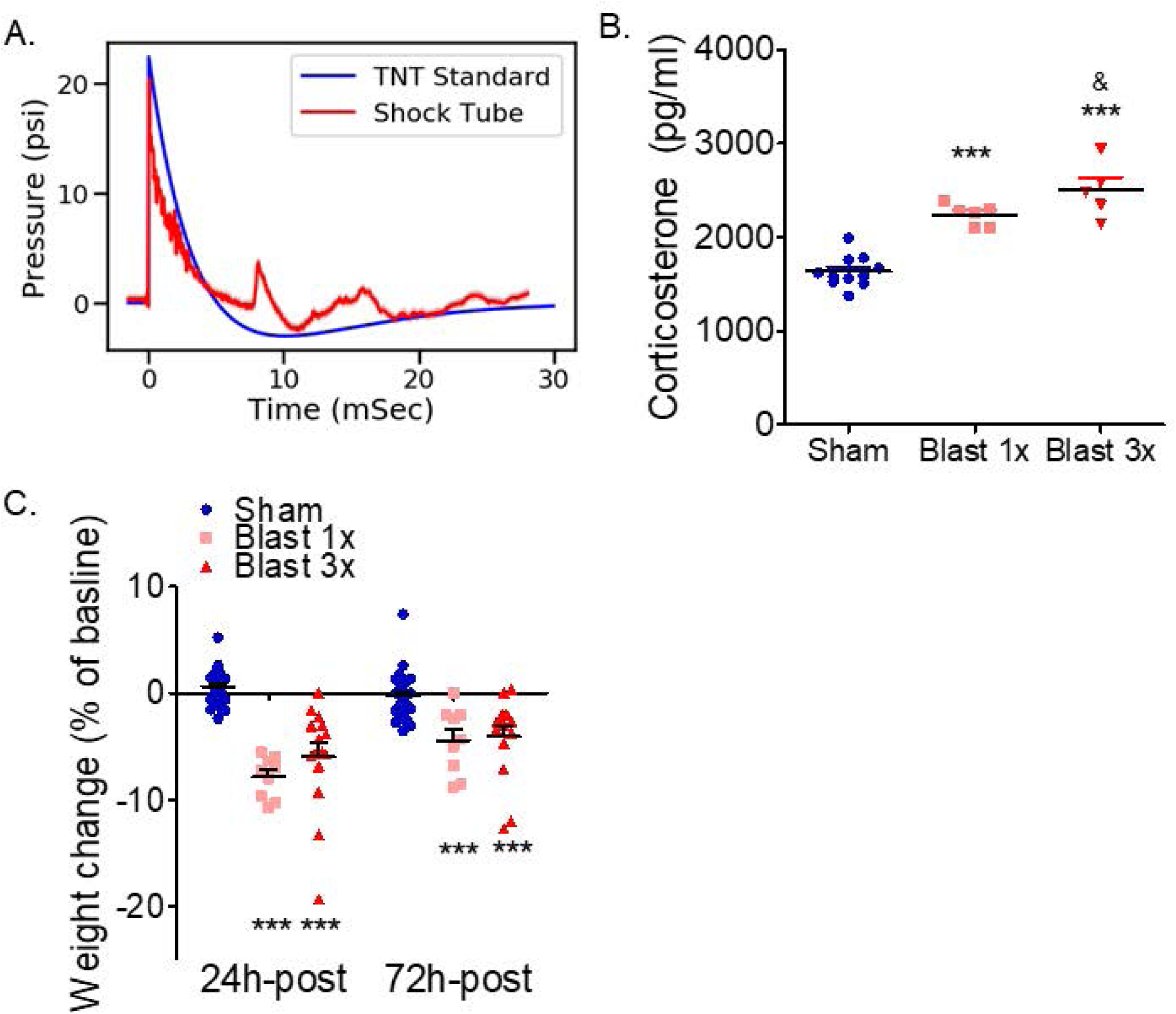
Blast-induced acute stress response. (a) Time versus pressure plot (averaged over 350 blasts) measuring the static blast overpressure (measured 5 cm above the animal). Note close correspondence with the superimposed Friedlander waveform expected from an open-field detonation of approximately 20 kg of TNT at a distance of 7.8 m. (b) Blast exposure results in blood corticosterone increase. \*\*\**p* ≤ 0.0001: sham vs. blast. (c) Blast exposure results in acute weight loss that lasts at least three days. \*\*\**p* ≤ 0.0001: sham vs. blast.

### Odorant conditioning paradigm

See Figure 2A for experimental schematic. One week prior to sham/blast exposure, animals were first pre-exposed to a Plexiglas T-Maze (66 cm long × 40 cm wide × 15 cm high) for 5 min. A stainless-steel mesh tea-ball (Amazon) containing a clean quarter Nestletts (PharmaServ, Framingham, MA) was also placed into each home cage to allow for pre-exposure. The following week, sham/blast exposure occurred on three consecutive days. On each day of sham/blast exposure, animals received a stainless-steel mesh tea-ball (Amazon) containing one-quarter Nestletts (PharmaServ, Framingham, MA) with 20 μl of imitation almond extract (Kroger, Cincinnati, OH) in their home cage five minutes prior to sham/blast exposure. The Nestletts and scent were refreshed on each morning of sham/blast exposure and the tea-ball remained in place until 24 hours following the final sham/blast exposure. One month following exposure, animals were tested for odorant-conditioning in the T-Maze with a tea-ball containing one-quarter Nestlett with 20 μl imitation almond odorant cue placed in the left arm of the maze and a tea-ball containing one-quarter Nestlett with 20 µl saline placed in the opposite arm of the maze. Animals were placed in the long arm of the T-maze and given 5 min to explore the entire maze. Latency to enter and time spent in each of the two distal ends of the short arms was recorded and analyzed using Anymaze (Stoelting, Wood Dale, IL). There were no statistically significant differences between 1x sham (n = 5) and 3x sham (n = 6)-treated mice (saline corner time: Student’s unpaired t-test, t[9]=0.537, *p*>0.05; odor corner time: Student’s unpaired t-test, t[9]=0.036, *p*>0.05; saline corner latency: Student’s unpaired t-test, t[9]=0.712, *p*>0.05; odor corner time: Student’s unpaired t-test, t[9]=1.113, *p*>0.05), thus, 1x and 3x sham animals were pooled together for subsequent odorant-pairing analyses.

**Figure 2:**
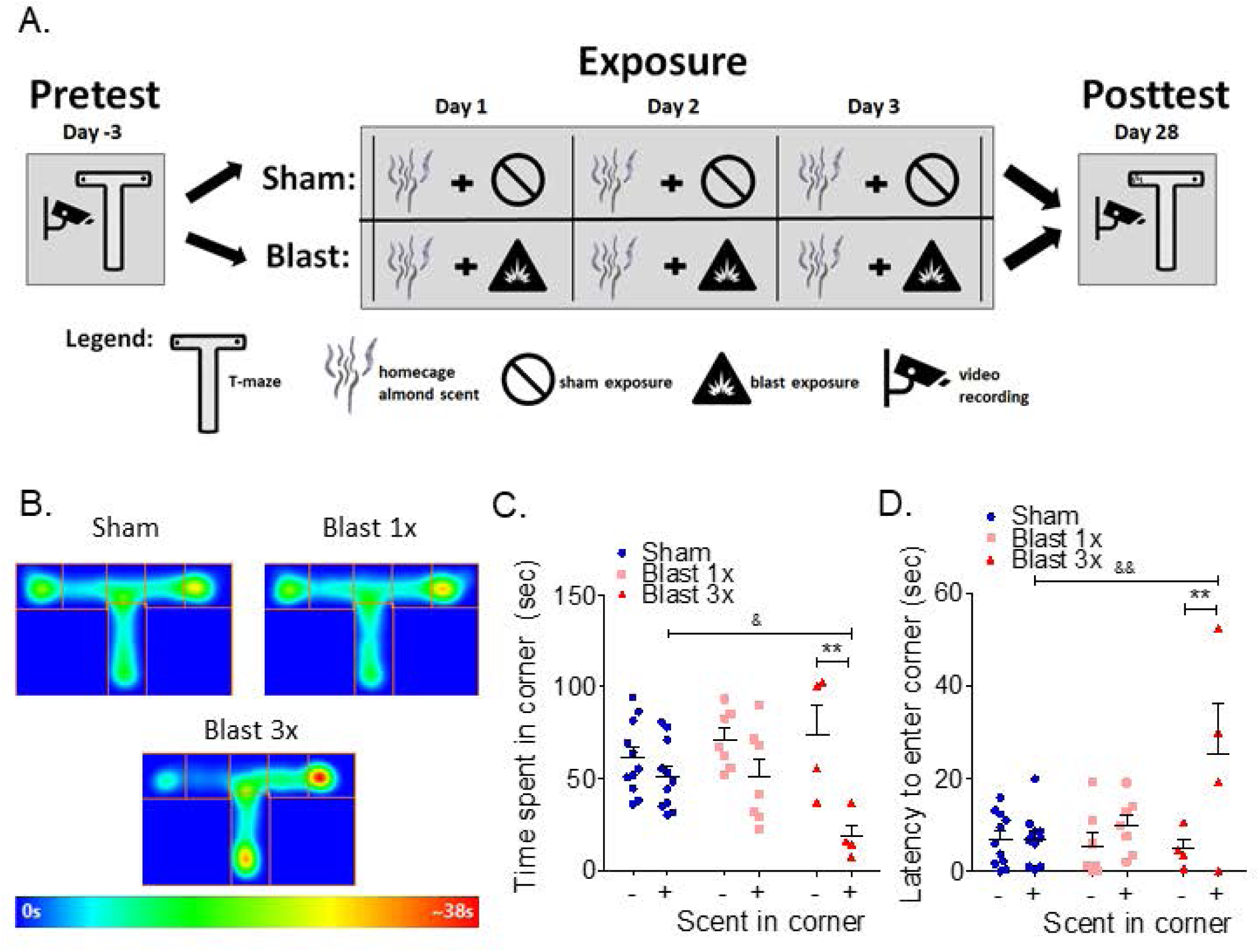
Blast-induced odorant aversion. (a) Schematic of odorant-blast aversion paradigm. (b) Heat maps of odorant aversion post-test (odorant placed in left corner of T). (c) Significant aversion to repetitive blast/odor pairings one-month post-injury, with no aversion seen from sham or a single blast/odor pairing. (d) Latency to enter to odorant corner is significantly increased one-month post repetitive blast/odor pairings. Two-way RM ANOVA *post hoc* Bonferroni Multiple Comparison Test. \**p* ≤ 0.05 and \*\**p* ≤ 0.001: + odor corner vs. – odor corner, ^&^*p* ≤ 0.05 and ^&&^*p* ≤ 0.001: sham vs blast odor corner. Error bars are mean +/- SEM.

### Place conditioning paradigm

Figure 3A illustrates the experimental design. A balanced-three compartment conditioning apparatus was used as described previously (17). One week prior to sham/blast exposure, animals were pre-tested by placing individual animals in the small central compartment and allowing exploration of the entire apparatus for 20 min. Time spent in each compartment was recorded and analyzed using Anymaze. Mice were randomly assigned an AM and PM box (either gray walls or vertical black and white strip walls). The following week, sham/blast exposure occurred on three consecutive days. On each day of exposure, in the morning, animals were first placed in their AM-pairing chamber containing distinct visual cues for 10 minutes and then were immediately given a sham exposure. In the afternoon, animals were placed in their PM-pairing chamber containing a different set of distinct visual cues for 10 minutes and then were immediately given a blast or sham exposure (depending on group assignment). Place conditioning was assessed at one and three months following repetitive exposure by allowing the mice to roam freely in all three compartments. Time spent in each compartment was recorded and analyzed using Anymaze (Stoelting, Wood Dale, IL). Place conditioning scores were calculated by subtracting the time spent in the PM paired compartment from the time spent in the AM paired compartment.

**Figure 3:**
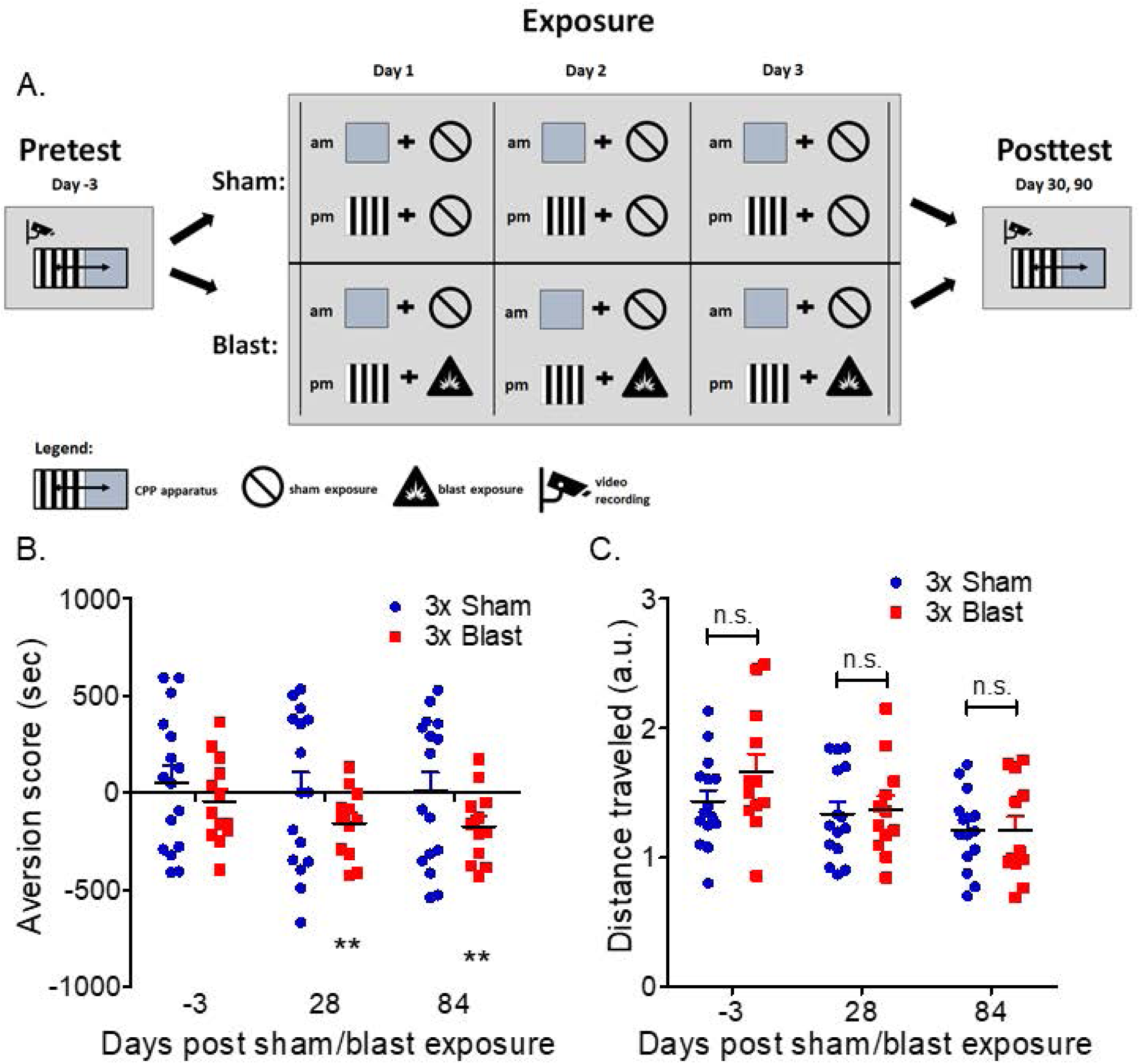
Blast-induced place aversion. (a) Schematic of blast-induced place aversion paradigm. (b) Significant aversion to blast-paired but not sham-paired cues at one month and three months post-injury. (c) No difference in locomotion during place aversion post-test across groups and timepoints. Student’s one-sample t-test vs. theoretical of 0 (no aversion). Two-way ANOVA *post hoc* Bonferroni. \**p* ≤ 0.05, \*\**p* ≤ 0.001, and \*\*\**p* ≤ 0.0001: sham vs blast or neutral vs. paired. Values represent mean ± SEM.

### Cue conditioning and re-exposure paradigm

See Figure 4A for experimental schematic. Sham/blast exposure occurred on three consecutive days. On each day of exposure, animals were first placed in a pairing chamber containing distinct visual cues (randomly assigned to gray or black and white striped walls) for 10 minutes and then were immediately given a sham or blast exposure (depending on group assignment). One month following repetitive exposure, the animals were re-exposed to either a neutral chamber or the chamber previously paired with blast or sham for 10 minutes, and movement (via Anymaze (Stoelting, Wood Dale, IL)) and ultrasonic vocalizations were recorded. Blood was collected from the submandibular vein one day prior and 30 minutes after removal from the pairing chamber. Plasma samples were processed to assay corticosterone levels using an ELISA kit as per manufacture protocol (Arbor Assays, Ann Arbor, MI). USVs were recorded using a Petterson microphone (Norway, model M500-384) and Avisoft SASLab Lite recording software and were manually analyzed using RavenLite (Cornell lab of Ornithology).

**Figure 4:**
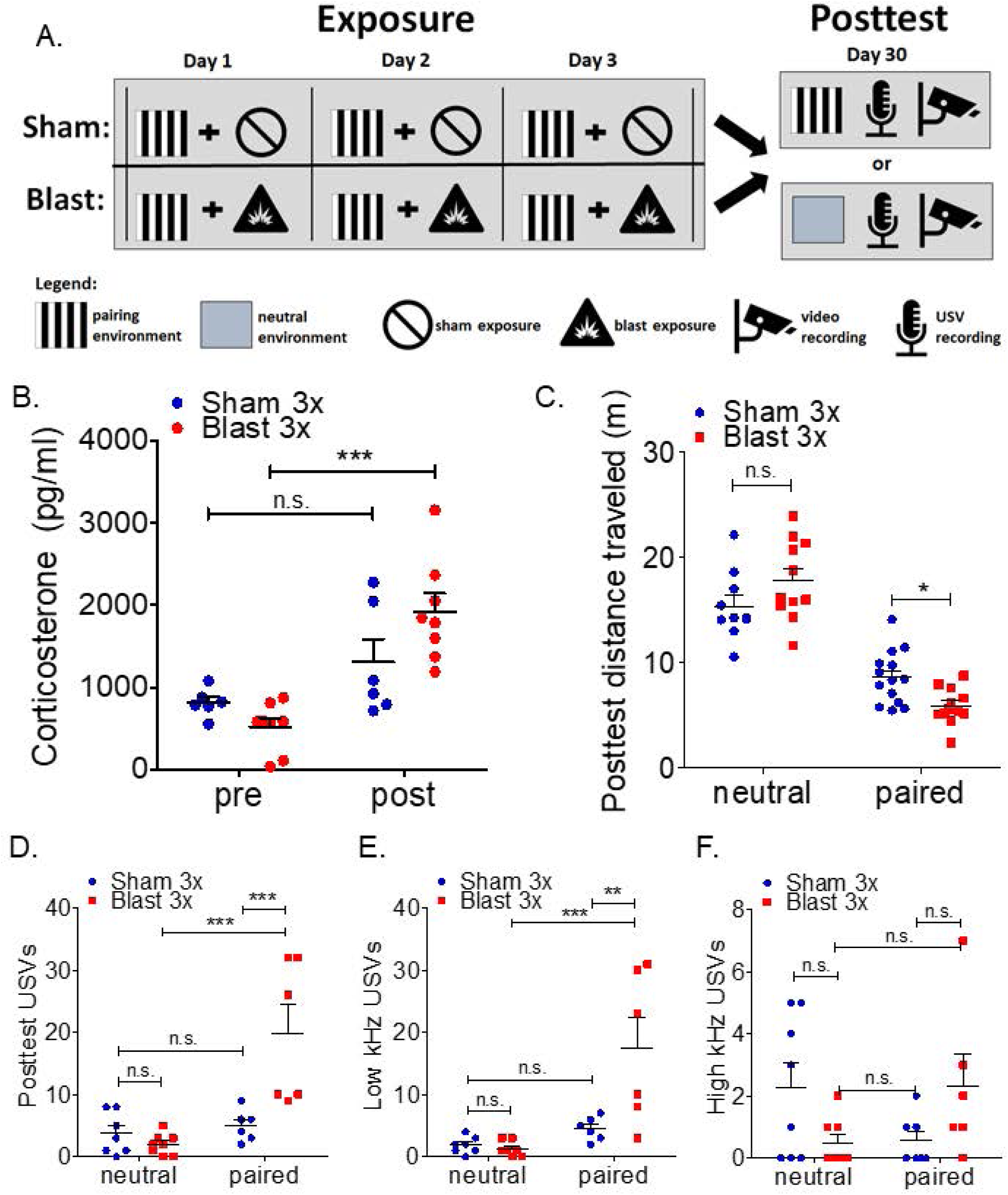
Blast-induced stress responses to cue re-exposure. (a) Schematic of blast-induced environmental pairing and re-exposure paradigm. (b) Significant increase in plasma corticosterone following re-exposure to blast-paired but not sham-paired cues one-month post-injury. (c) Significant decrease in locomotion during re-exposure to blast-paired but not sham-paired or neutral cues one-month post-injury. (d-f) Aversive (low kHz) ultrasonic vocalizations are significantly increased following re-exposure to blast-paired but not sham-paired or neutral cues one-month post-injury. Student’s one-sample t-test vs. hypothetical mean=1.0. Two-way ANOVA *post hoc* Bonferroni. \**p* ≤ 0.05, \*\**p* ≤ 0.001, and \*\*\**p* ≤ 0.0001: sham vs blast or neutral vs. paired. Values represent mean ± SEM.

### Data analysis

Where appropriate, data were analyzed using: (*i*) two-tailed Student’s t-tests; (*ii*) one-way or two-way (between/within subjects design) repeated or non-repeated measures analysis of variance (ANOVA), followed by Newman-Keuls Multiple Comparison Tests or Bonferroni Post-hoc tests, respectively. Reported *p* values denote two-tailed probabilities of p≤ 0.05 and non-significance (n.s.) indicates *p*> 0.05. Statistical analyses were conducted using Python, Graph Pad Prism 4.0 (GraphPad Software, Inc., La Jolla, CA), and SPSS software (IBM, Armonk, NY).

## RESULTS

### Acute stress responses following blast exposure

Using well-established methods (10, 11, 28-30), male C57Bl/6 adult mice were exposed to one or three blast overpressures (BOPs) using a pneumatic shock tube delivering a peak static pressure of 20.23 psi +/- 0.08 psi, positive phase duration of 5.80 ms +/- 0.02 ms; 0.037 psi*ms +/- 0.000 psi*ms (Figure 1A).

Stressful events commonly elicit release of corticosterone (in rodents), thus we measured the level of corticosterone in blood thirty minutes following exposure to one or three blast overpressures (or sham exposure). There were no statistically significant differences between 1x sham and 3x sham-treated mice. Thus, 1x and 3x sham animals were pooled together for subsequent corticosterone analysis (see Materials and Methods for sham comparison statistics). In accordance with an acute stress response, blast exposure resulted in a significant increase in blood corticosterone (one-way ANOVA: F[2,21]=41.79, p<0.0001, Bonferroni’s Multiple Comparison Test post-hoc: sham n=11, blast 1x n=6, blast 3x n=5). Post-hoc analyses demonstrate significant corticosterone increases in both 1x and 3x blast animals as compared to sham control. Likewise, there was a significant increase in corticosterone in 3x blast animals as compared to 1x blast animals (Figure 1B). Stress exposure in mice commonly results in weight loss, thus we measured body weight at baseline, 24h, and 72h following sham or blast exposure. There were no statistically significant differences between 1x sham and 3x sham-treated mice. Thus, 1x and 3x sham animals at each time point were pooled together for subsequent weight analyses (see Materials and Methods for sham comparison statistics). In accordance with an acute stress response, blast exposure resulted in acute weight loss (two-way RM ANOVA: interaction effect F[2,46]=12.95, p<0.0001, Bonferroni’s Multiple Comparison Test post-hoc: sham n=25, blast 1x n=9, blast 3x n=15). Post-hoc analyses demonstrate significant weight loss for both 1x and 3x blast animals at both 24h and 72h following exposure (Figure 1C).

### Chronic aversion to cues previously paired with blast

To further investigate potential blast-induced stress responses, we employed a conditioned odorant paradigm in mice that pairs sham or blast exposure with a neutral odorant cue (Fig. 2A). The neutral odorant cue (almond scent) was placed in the home cage five minutes prior to the first sham or blast exposure and remained in place (scent refreshed daily) until 24 hours post final sham or blast exposure. Finally, conditioning to the paired-odorant cue was assessed one month following exposure. There were no statistically significant differences between 1x sham and 3x sham-treated mice. Thus, 1x and 3x sham animals at each time point were pooled together for subsequent odorant pairing analyses (see Materials and Methods for sham comparison statistics). The panels in Figure 2B show occupancy heat maps for each group during the post-test (odorant is placed in left corner of T-maze) one month following the exposure/odorant pairings. Mice developed a significant aversion to the odorant when subsequently presented alone in one corner of the testing chamber (left corner of T-maze) (two-way RM ANOVA: main effect of location F[1,19]=11.83, p=0.003, Bonferroni’s Multiple Comparison Test post-hoc: sham n=11, blast 1x n=7, blast 3x n=4) (Figure 2C). Post-hoc analyses demonstrated significant aversion following only repetitive (3x, one per day) but not single (1x) blast or sham exposure. Odorant-blast pairings also increased the latency to enter the odorant corner in a blast number-dependent manner (two-way RM ANOVA: main effect of location F[1,19]=8.798, p=0.008, Bonferroni’s Multiple Comparison Test post-hoc: sham n=11, blast 1x n=7, blast 3x n=4) (Figure 2D).

To assess the generality of these observations, the ability of repetitive blast exposure to induce place aversion was investigated using a modified place conditioning paradigm in mice that pairs sham and blast exposure with distinct visual cues (Fig. 3A). On each day of exposure, in the morning, animals were first placed in a pairing chamber containing distinct visual cues for 10 minutes and then were immediately given a sham exposure (e.g. anesthesia only), then in the afternoon, animals were placed in a pairing chamber containing a different set of distinct visual cues for 10 minutes and then were immediately given a blast or sham exposure. Conditioned aversion to the blast-paired compartment was evident one month post-blast exposure and was sustained until at least three months post-blast exposure (one-sample t-test vs. a theoretical mean of 0 (no aversion):; sham baseline: t[15]=1.372, p=0.1903, sham 1 month): t[14]=1.479, p=0.1613, sham 3 months: t[13]=0.5922, p=0.5639; blast baseline: t[14]=0.1919, p=0.8506 blast 1 month: t[14]=3.346, p=0.0048, blast 3 months: t[12]=3.256, p=0.00679; (Figure 3B). Despite significant place aversion, blast exposure did not affect total distance traveled during the post-test, suggestion that aversion was not an nonspecific effect of locomotion changes (two-way RM ANOVA: interaction F[2,50]=0.931, p=0.401, Bonferroni’s Multiple Comparison Test post-hoc: sham n=16, blast 3x n=15) (Figure 3C). Taken together, these data demonstrate that a single blast exposure was sufficient to produce acute stress responses. However, only repetitive blast exposure induced significant conditioned aversion.

### Dysphoric and aversive behavioral response to visual cues previously paired with blast

In order to investigate whether cues previously paired with blast or sham exposure elicit distinct physiological and behavioral responses during re-exposure, we extended the modified place conditioning paradigm used above to allow for re-exposure to either a neutral visual environment or one that was previously paired with exposure (Figure 4A). On each day of exposure, animals were first placed in a pairing chamber containing distinct visual cues for 10 minutes and then were immediately given a sham or blast exposure. One month following repetitive exposure, animals were re-exposed to either a neutral chamber or the chamber previously paired with exposure (blast or sham) for 10 minutes, movement and ultrasonic vocalizations (USVs) were recorded, and blood was collected (pre/post re-exposure) for subsequent plasma corticosterone analysis. Differential blood corticosterone responses were detected pre/post re-exposure to sham/blast-paired cues (two-way ANOVA: interaction F[1,12]=7.998, p=0.0152, Bonferroni’s Multiple Comparison Test post-hoc: sham n=6, blast 3x n=8) (Figure 4B). Post-hoc analyses demonstrate a significant corticosterone increase in blast but not sham animals (pre/post re-exposure). Differential behavioral responses were also demonstrated following re-exposure to neutral vs. sham/blast-paired cues (two-way ANOVA: interaction effect F[1,43]=9.462, p=0.0036, Bonferroni’s Multiple Comparison Test post-hoc: sham n=13, blast 3x n=11) (Figure 4C). Post-hoc analyses demonstrated a significant decrease in distance traveled during re-exposure to blast-paired cues as compared to sham-paired cues with no difference when re-exposed to neutral cues. In addition to movement, we also recorded ultrasonic vocalizations (USVs) during re-exposure as an additional read-out of affective state. Total USVs were significantly increased in blast-exposed mice during re-exposure to the sham/blast-paired but not neutral-paired cues (two-way ANOVA: interaction effect F[1,23]=9.714, p=0.0049, Bonferroni’s Multiple Comparison Test post-hoc: sham n=8, blast 3x n=6) (Figure 4D). In line with an aversive/dysphoric stress response to reminders of blast exposure, low kHz USVs were specifically increased during re-exposure to the sham/blast paired but not neutral-paired cues (two-way ANOVA: interaction effect F[1,23]=8.820, p=0.0069, Bonferroni’s Multiple Comparison Test post-hoc: sham n=8, blast 3x n=6) (Figure 4E). Post-hoc analyses demonstrated a significant increase in low kHz USVs during re-exposure to blast-paired cues as compared to sham-paired cues with no difference when re-exposed to neutral cues. While a significant interaction effect was also seen in the number of high kHz USVs (two-way ANOVA: interaction effect F[1,23]=5.206, p=0.0321, Bonferroni’s Multiple Comparison Test post-hoc: sham n=8, blast 3x n=6) (Figure 4F), post-hoc analyses did not demonstrate significant differences between cues or blast and sham exposure.

## DISCUSSION

Blast-induced mTBI is currently defined as cellular and/or structural damage to the brain, which can cause adverse somatic, vestibular, cognitive, and affective symptoms (2, 33). PTSD is an anxiety disorder which develops after exposure to potentially life-threatening stress (such as blast) and persistent symptoms include re-experiencing, avoidance, and hyperarousal (21, 34, 35). As such, blast-induced mTBI is commonly associated with the development of battlefield PTSD and the two conditions share several overlapping symptoms. Indeed, some have postulated that postconcussive symptoms following mTBI are non-specifically related to physical blast injury, and instead better explained by psychological trauma consistent with PTSD (6, 12). Because battlefield blast exposures can simultaneously provoke both psychological and neurological insults, understanding the basis for chronic symptoms in dual-diagnosis (e.g. mTBI and PTSD) Veterans has remained challenging.

Translational research efforts using rodent models are thought to be free from many of the confounding variables associated with human exposures in the battlefield, providing much needed insight into underlying mechanisms by which blast exposure produces dysfunction, but only in so far as these models are sufficiently translationally relevant. Drawing from long-established chronic stress and Pavlovian conditioning research (16-22), the results herein strongly argue that the conditions surrounding the immediate blast exposures per se, are sufficient to drive Pavlovian learning and generate subsequent PTSD-like behavioral outcomes. Specifically, these data directly support the notion that blast exposure in this animal model can give rise to stress responses (both physiological and behavioral) that elicit subsequent PTSD-like avoidance, intrusive symptoms, and mood alteration (e.g. dysphoria) (36, 37). We utilized place conditioning as an objective measure of behavior and to infer a relationship between the aversion exhibited at time of testing to previous stress and dysphoria at the time of conditioning. Indeed, Land et al., (2008) operationally defined dysphoria as the emotional response to a sustained stimulus that creates aversion and is thought to represent the underlying emotional state at time of stimulus exposure. Importantly, we demonstrate chronic effects which persist at least three months following blast exposure, an important requirement to differentiate PTSD from acute stress disorder in humans as per the Diagnostic and Statistical Manual of the American Psychological Association (DSM). While PTSD-like aversion-related outcomes were only seen in animals exposed to repetitive blast, acute stress effects related to increased corticosterone and weight loss were apparent following a single blast exposure, raising the possibility that a single blast might model aspects of acute stress disorder without progressing to full PTSD-like outcomes.

In most animal studies, blast exposure is carried out under anesthetized, thus potentially restricting opportunities for the animals to form cognitive associations with the immediate blast exposure. Under such conditions it is well-established that blast exposure produces chronic PTSD-like behavioral responses such as increased anxiety and exaggerated startle response, raising the possibility that cellular/structural insults to the brain caused by blast may be sufficient to drive pathogenic processes leading to PTSD-like symptoms with limited appeal to psychological trauma and/or stress (15). Some studies have used an additional stressor at the time of injury (e.g. dual exposure) in order to study mTBI/PTSD comorbidity (38-41). The findings in this report demonstrate that repetitive blast exposure alone can generate traumatic stress responses sufficient to drive subsequent aversion and dysphoria, despite anesthesia and without the requirement for an additional, experimentally administered stressor.

In a typical pre-clinical Pavlovian fear/stress conditioning experiment, a rodent receives repeated presentations of a conditioning stimulus (CS, e.g. an environmental chamber, odor, or auditory cue) that coincide with presentation of an unconditioned stimulus (US, e.g. shock, predator odor, or social defeat). Subsequently, the animal will display a variety of conditioned responses upon later re-exposure to the CS (e.g. avoidance, freezing, increased heart rate and blood pressure, corticosterone release, ultrasonic vocalizations). Importantly, the typical blast exposure paradigm provides opportunities for Pavlovian learning (even in the absence of experimenter-delivered conditioned stimuli). For example, animals must first be transported from the vivarium to a blast holding room where they stay until transferred to the blast exposure room. Once transferred to the blast exposure room, animals are placed in an anesthesia induction chamber, anesthetized, exposed to blast, and then allowed to recover. Overall, while the actual blast exposure under anesthesia might only last 3-5 minutes, the entire episode outside of the vivarium might last anywhere from 10 minutes to hours (depending on location of vivarium vs. blast holding and exposure rooms). In the current study we exposed animals to discrete Pavlovian cues (i.e. an almond scent or a distinct visual environment), but we also postulate that additional cues available surrounding events such as transport to and wait time in the blast holding room, transport to the blast exposure room, and placement in the anesthesia induction chamber might also serve as potential conditioning stimuli without intentional experimenter administration or manipulation. As such, longstanding cognition and behavior literature supports the idea that being awake and conscious during a traumatic event is not required for subsequent aversion and PTSD-like behavioral outcomes (e.g. drug-facilitated sexual assault, post-intensive care syndrome, and conditioned taste aversion to anesthetics (42-45). Indeed, we take precautions to limit unnecessary re-exposure to potential blast-related cues (for example, we use different colored drapes and gowns on blast exposure days vs. behavioral testing days, we do not use the holding or blast rooms for any other procedures).

The idea that sedation can be used to dissect psychological trauma from neurological and cellular injury is inherently attractive from a reductionist viewpoint, especially when considering the contemporary significance of an as yet, unresolved controversy regarding the underlying causes of the “shell shock” symptoms first described during World War I (46, 47). The data herein demonstrate that repetitive blast is capable of provoking PTSD-like behavioral outcomes related to aversion and dysphoria, as well as the traumatic brain insults we and others have reported previously in this mouse model (10-15). Thus, on one hand this suggests that animal blast models can, like veterans with blast-related mTBI/PTSD, present a more complex biological puzzle than one might hope for. On the other hand, and perhaps more importantly, the findings in this report are encouraging because they demonstrate a previously unappreciated level of translational relevance in this animal model. Indeed, using blast-paired Pavlovian cues, it may be possible to carry out increasingly relevant studies of trauma re-exposure effects on mTBI- and PTSD-related adverse outcomes such as sleep quality, substance use and misuse, and aggression, facilitating understanding of the mechanisms of blast-induced mTBI/PTSD like outcomes and speeding progress toward better treatments for Veterans and servicemembers with long-term blast-related health concerns.

## Acknowledgments

This work was supported by a Department of Veteran Affairs (VA) Basic Laboratory Research and Development (BLR&D) Career Development Award 1IK2BX003258 (AGS), a VA BLR&D Merit Review Award 5I01BX002311 (DGC), a VA CSR&D Career Development Award #CX-001787 (GET), VA Rehabilitation Research and Development Service Merit Review Award #B77421 (ERP), University of Washington Friends of Alzheimer’s Research (DGC, ERP), UW Royalty Research Fund (DGC), and the VA Northwest Mental Illness Research Education and Clinical Center (JSM, MAR, ERP). We thank Greg Elder for insightful comments on the manuscript and would like to thank Scott Ng-Evans, Benjamin Land, Traci J Webber, Cindy Pekow, Kari Koszdin, and Monica Foley for considerable technical assistance and veterinary care.

## Disclaimer

The views expressed in this scientific presentation are those of the author(s) and do not reflect the official policy or position of the U.S. government or the Department of Veterans Affairs.

